# Neutralization and Stability of SARS-CoV-2 Omicron Variant

**DOI:** 10.1101/2021.12.16.472934

**Authors:** Cong Zeng, John P. Evans, Panke Qu, Julia Faraone, Yi-Min Zheng, Claire Carlin, Joseph S. Bednash, Tongqing Zhou, Gerard Lozanski, Rama Mallampalli, Linda J. Saif, Eugene M. Oltz, Peter Mohler, Kai Xu, Richard J. Gumina, Shan-Lu Liu

**Author notes:** Authors Contributed Equally to this Work (Cong Zeng, John P. Evans, Panke Qu).

## Abstract

The SARS-CoV-2 B.1.1.529/Omicron variant was first characterized in South Africa and was swiftly designated a variant of concern^1^. Of great concern is its high number of mutations, including 30-40 mutations in the virus spike (S) protein compared to 7-10 for other variants. Some of these mutations have been shown to enhance escape from vaccine-induced immunity, while others remain uncharacterized. Additionally, reports of increasing frequencies of the Omicron variant may indicate a higher rate of transmission compared to other variants. However, the transmissibility of Omicron and its degree of resistance to vaccine-induced immunity remain unclear. Here we show that Omicron exhibits significant immune evasion compared to other variants, but antibody neutralization is largely restored by mRNA vaccine booster doses. Additionally, the Omicron spike exhibits reduced receptor binding, cell-cell fusion, S1 subunit shedding, but increased cell-to-cell transmission, and homology modeling indicates a more stable closed S structure. These findings suggest dual immune evasion strategies for Omicron, due to altered epitopes and reduced exposure of the S receptor binding domain, coupled with enhanced transmissibility due to enhanced S protein stability. These results highlight the importance of booster vaccine doses for maintaining protection against the Omicron variant, and provide mechanistic insight into the altered functionality of the Omicron spike protein.

---

Since the introduction of SARS-CoV-2 into the human population in late 2019, adaptive evolution of the virus has resulted in increased transmissibility and resistance to vaccine- or infection-induced neutralizing antibodies^2,3^. Indeed, the initial D614G mutation in the virus spike (S) protein enhanced virus stability, infectivity, and transmission^4,5^. This initial adaptation was followed by the emergence of several SARS-CoV-2 variants of concern, including Alpha (B.1.1.7), which rapidly spread from Europe to become the global dominant variant in other parts of the world^6^. Subsequently, the Beta (B.1.351) variant exhibited substantial resistance to neutralization^7^, although failed to disseminate as widely. The most recent wave of the pandemic is attributed to the Delta (B.1.617.2) variant characterized by moderate neutralization resistance combined with enhanced transmissibility, driving its dominance worldwide^8^. Despite these evolutionary leaps, vaccine-mediated protection from severe disease and hospitalization remained high^9^.

The emergence of the Omicron (B.1.1.529) variant has generated serious concern about the continued efficacy of vaccines and the future course of the pandemic^10,11^. In addition to its sheer number of mutations^12^, Omicron contains specific alterations that have previously been shown to impact vaccine resistance, namely in the receptor-binding domain (RBD), a primary target of host neutralizing antibodies to the S protein, as well as a number of other non-RBD mutations, including some in the S2 subunit (**Fig. 1a**). Moreover, when present in a geographic region, the Omicron variant constitutes a rapidly increasing proportion of COVID-19 cases^13,14^, suggesting a further enhancement of transmissibility.

**Figure 1.**
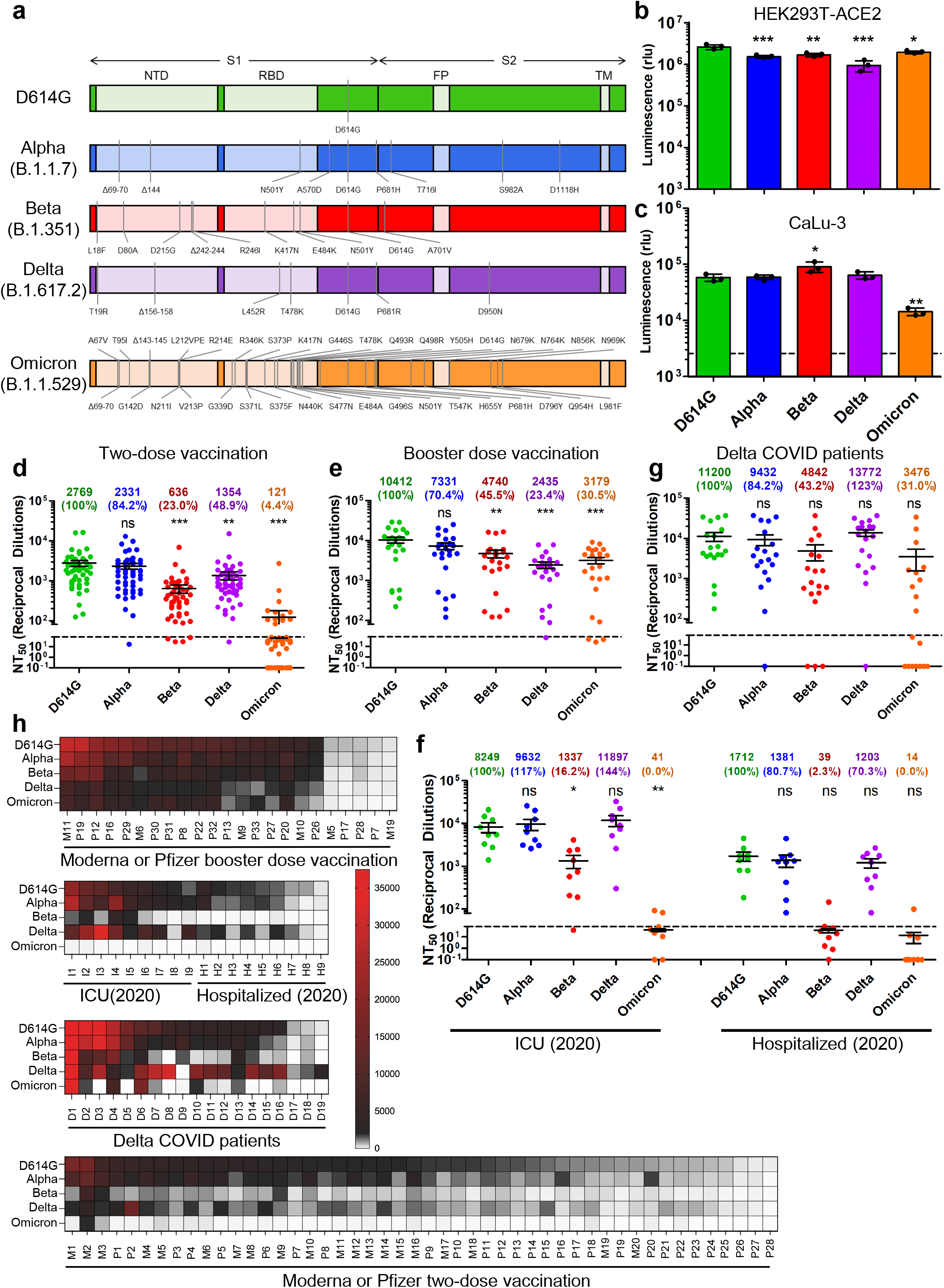
The Omicron SARS-CoV-2 variant exhibits strong immune-escape that is overcome by booster vaccination. (**a**) Diagrams of SARS-CoV-2 S variants used for pseudotyping, which indicate the location of specific mutations as well as the S1 and S2 subunits of S, the N-Terminal Domain (NTD), Receptor Binding Domain (RBD), Fusion Peptide (FP), and Trans-membrane domain (TM). (**b**) Infectivity of pseudotyped viruses produced in parallel for infection of HEK293T cells stably expressing ACE2. (**c**) Infectivity of pseudotyped lentivirus in human lung epithelia-derived CaLu-3 cell line. Bars in b and c represent means +/- standard deviation, and significance is determined by one-way ANOVA with Bonferroni’s multiple testing correction. Results of at least 3 independent experiments are averaged and shown. (**d**) Sera from 48 HCWs collected 3-4 weeks after second mRNA vaccine dose was used to neutralize pseudotyped virus for variants, and the resulting 50% neutralization titers (NT_50_) are displayed. (**e**) Sera from 23 HCWs following homologous mRNA booster vaccination were assessed for nAb titers. (**f**) Sera from 9 ICU COVID-19 patient samples and 9 hospitalized non-ICU COVID-19 patient samples collected in 2020 prior to the approval of any SARS-CoV-2 vaccines were assessed for nAb titers. (**g**) Sera from 19 ICU COVID-19 patient samples collected during the Delta-wave of the pandemic were assessed for nAb titers. Mean NT_50_ values in panels d-g are displayed at the top of plots along with relative neutralization sensitivity with D614G set to 100%; bars represent mean +/- standard error, and significance relative to D614G is determined by one-way ANOVA with Bonferroni’s multiple testing correction. (**h**) Heat maps showing patient/vaccinee NT_50_ values against each variant. Patient/vaccinee numbers are identified as P for Pfizer/BioNTech BNT162b2 vaccinated/boosted HCW, M for Moderna mRNA-1273 vaccinated/boosted HCW, I for ICU patient samples collected during the 2020 D614G-wave, H for hospitalized non-ICU patient samples collected during the 2020 D614G-wave, or D for ICU patient samples collected during the Delta-wave. P-values are represented as *p < 0.05, **p < 0.01, ***p < 0.001; ns, not significant.

## Resistance of Omicron to neutralizing antibodies in two-dose vaccinees, booster-dose recipients and COVID-19 patients

Given heightened concerns about Omicron’s transmissibility and vaccine resistance, we examined the its S protein compared to other major SARS-CoV-2 variants using a previously reported pseudotyped lentivirus system^15^. We first examined infectivity using lentivirus pseudotypes and HEK293T-ACE2 as target cells. As shown in **Fig. 1b**, the infectivity of Omicron was largely comparable to the other major variants, all of which were lower than the ancestral D614G. However, in a more physiologically relevant host, human lung epithelia-derived CaLu-3 cells, Omicron exhibited reduced infectivity relative to D614G (**Fig. 1c**).

We next examined the ability of Omicron to escape vaccine-induced neutralizing antibodies, a critical measure of protection from SARS-CoV-2 infection^16^. To address this, we collected sera from 48 health care workers (HCWs) 3-4 weeks post-second dose of either Moderna mRNA-1273 (n = 20) or Pfizer/BioNTech BNT162b2 (n = 28). Having previously examined this cohort for the ability of the D614G, Alpha, Beta, and Delta variants to escape serum antibody neutralization (nAb)^17^, we compared the neutralization resistance of Omicron to these variants of concern. We found that the Omicron variant exhibited significantly more neutralization resistance, i.e., 22.9-fold (p < 0.001), compared to ancestral D614G, with the Alpha, Beta, and Delta variants exhibiting a 1.2-fold, 4.4-fold (p < 0.001), and 2.0-fold (p < 0.01) decrease in nAb titers, respectively (**Fig. 1d**). In total, only 27.1% (13/48) of HCWs exhibited nAb titers against Omicron above the detection limit (NT_50_ < 80); however, several individuals (1-3) exhibited strong nAb titers that were maintained against Omicron (**Fig. 1d and h)**. Moderna mRNA-1273 in HCWs slightly outperformed Pfizer/BioNTech BNT162b2 (**Fig. S1a**), as we have reported previously ^17,18^.

We further sought to examine resistance to vaccine-induced immunity for Omicron along with D614G, Alpha, Beta, and Delta for 23 HCWs 1-11 weeks post-booster vaccination. We found that booster vaccination not only increased the nAb NT_50_ titer against all variants, including D614G, but also significantly restored neutralization of Omicron, with only 3.3-fold reduced NT_50_ relative to D614G, which was even less than the 4.3-fold reduction for the Delta variant (**Fig. 1e**). These data indicate that booster dose administration not only enhances nAb titers, but also enhances the breadth of the nAb response, especially against Omicron. For 18 HCWs, we analyzed the post-second dose and post-booster dose samples, which showed significantly higher nAb titers following booster vaccination (**Fig. S1b-f**). Pfizer/BioNTech BNT162b2-vaccinated HCWs exhibited slightly higher nAb titers than Moderna mRNA-1273-vaccinated HCWs post-booster (**Fig. S1g**).

We further examined nAb resistance of Omicon and other major variants in ICU (n = 9) and hospitalized non-ICU (n = 9) patient serum samples collected during the 2020/D614G wave of the pandemic prior to vaccination. We found that Omicron was completely resistant to D614G-wave patient serum samples, with only 22.2% (2/9) of ICU and 11.1% (1/9) of hospitalized patients exhibiting a threshold of nAb titers (**Fig. 1f and h**). We further examined ICU patient samples (n = 19) collected during the Delta wave of the pandemic, including 5 confirmed as Delta infections by sequencing of virus from nasal swabs. Serum from these infected patients exhibited potent neutralization of Delta, as would be expected, while Omicron still exhibited strong resistance (**Fig. 1g and h**). Although mean titers from these individuals were comparable to boosted HCWs, a more significant proportion, 47.4% (9/19), exhibited no detectable nAb titers against Omicron (**Fig. 1g and h**). Further, this population contained 5 patients fully vaccinated with two doses, 1 fully vaccinated with 1 Johnson & Johnson dose, and 1 partially vaccinated (1 dose of Pfizer), which dramatically out-performed unvaccinated patients against Omicron and other variants, except Delta (**Fig. S1h**).

## Omicron spike exhibits reduced ACE2 binding, furin cleavage and cell-cell fusion

To better understand the impact of the Omicron mutations on binding to the virus receptor ACE2, we transfected HEK293T cells with variant spike constructs and determined S surface expression, as well as their capacity to bind to soluble ACE2 (sACE2), using flow cytometry (**Fig. S2a-d**). We observed comparable levels of S expression for Alpha, Beta, and Delta relative to D614G, with Omicron showing a slight reduction (**Fig. 2a, Fig. S2a-c and e**), as detected by a polyclonal antibody T62 against S1 or by anti-FLAG antibody specific for the N-termini of each recombinant spike. However, by normalizing for S surface expression in five independent experiments, we observed that Omicron variant exhibited a 2.4-fold reduced binding to sACE2 compared to the ancestral D614G (**Fig. 2b-c, Fig. S2d-f**).

**Figure 2.**
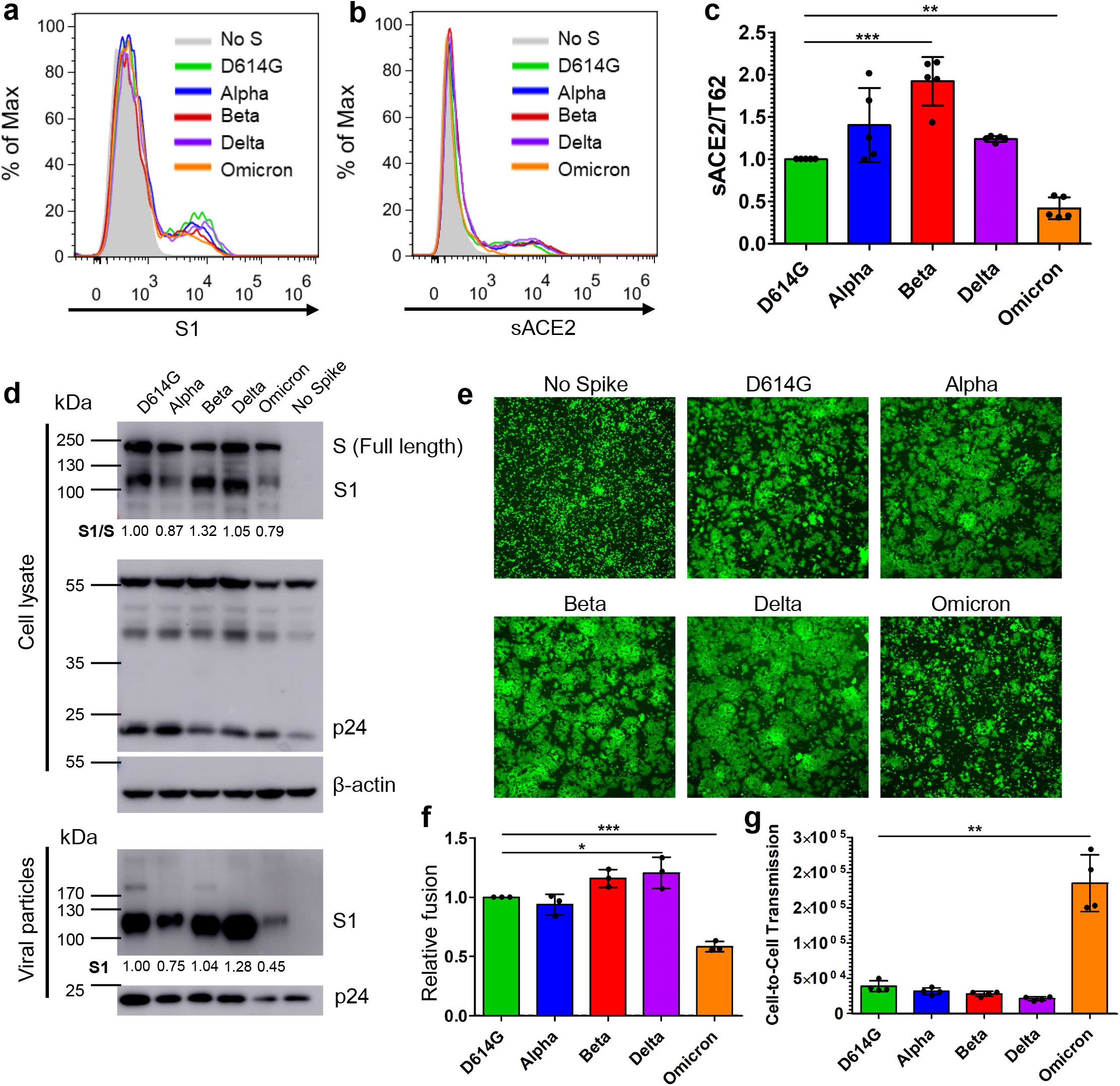
The Omicron S exhibits reduced ACE2 binding, processing, and fusion, but enhanced cell-to-cell transmission. (**a-c**) HEK293T cells were transfected with S expression constructs and stained for FACS with anti-S1 (T62) (**a**) or soluble ACE2-Fc fusion protein (sACE2) (**b**). (**c**) Mean fluorescence intensity for sACE2 signal was normalized with S1 signal to determine relative ACE2 binding; n=5. (**d**) Pseudotyped virus producer cells were lysed and pseudotyped virus was purified by ultracentrifugation and blots were probed for S1 subunit, lentivirus capsid (p24), and β-actin loading control; S cleavage was quantified using NIH ImageJ and by setting the ratio of S1/S of D614G to 1.00. (**e**) HEK293T cells transfected with GFP and S constructs were co-cultured with HEK293T-ACE2 cells for 24 hrs and imaged to visualize cell-cell fusion. (**f**) HEK293T cells transfected with S constructs and Tet-Off were co-cultured with HEK293FT-mCAT-Gluc cells transfected with an ACE2-GFP plasmid, and fusion-induced *Gaussia* luciferase signal was assessed 24 hrs after co-culture. (**g**) HEK293T cells transfected with S constructs to pseudotype *Gaussia* luciferase bearing lentivirus and were co-cultured with HEK293T-ACE2 cells and secreted *Gaussia* luciferase was assayed 24 hrs after co-culture. Bars in panels c, f and g represent means +/- standard deviation, and significance was determined by one-way ANOVA with Bonferroni’s multiple testing correction. Results were from at least three independent experiments. P-values are represented as *p < 0.05, **p < 0.01, ***p < 0.001.

We further sought to examine the processing and incorporation of the Omicron S protein into viral particles. Pseudotyped virus was purified and probed for the S1 subunit of S alongside virus producer cell lysate. We found that Omicron S, similar to that of Alpha, exhibited a reduced furin processing into S1 compared to D614G and Beta; consistent with prior reports^8,19^, the Delta variant showed increased furin cleavage in the spike (**Fig. 2d**). Notably, substantially less S1 was detectable in purified Omicron pseudotyped virus (**Fig. 2d**), despite a modestly low level of p24, which may reflect reduced incorporation of the Omicron S into pseudotyped virions and accounts for its decreased infectivity compared to D614G.

We next examined the efficiency of Omicron S protein-mediated cell-cell fusion. HEK293T cells were transfected with GFP and S protein constructs, and then co-cultured with HEK293T cells stably expressing ACE2 (HEK293T-ACE2) for 24 hrs. Cell-cell fusion was imaged using fluorescence microscope. We observed a marked reduction in the size of syncytia induced by Omicron S compared to other variants, especially by Delta and Beta (**Fig. 2e**). These results were confirmed using a more quantitative Tet-Off-based fusion assay, where spike and Tet-Off-expressing HEK293T cells were co-cultured with HEK293FT-mCAT-Gluc cells expressing ACE2. Upon cell-cell fusion, Tet-Off would induce the expression of *Gaussia* luciferase driven by a tetracycline-controlled transcription factor^20^. The Omicron variant exhibited 1.7-fold lower cell-cell fusion compared to D614G (p < 0.001) (**Fig. 2f**), consistent with reduced efficiency in mediating fusion at the plasma membrane.

## Omicron spike has enhanced capacity for cell-to-cell transmission

We reported the ability of SARS-CoV-2 spike to mediate cell-to-cell transmission^20^, To measure the efficiency of this process by Omicron S, we co-cultured with HEK293T-ACE2 target cells with HEK293T pseudotyping virus-producer cells and assessed cell-to-cell transmission after 24 hrs. Unexpectedly, we found that Omicron drastically increased the efficiency of cell-to-cell transmission, with 4.8-fold higher levels than D614G and other variants (**Fig. 2g**);, despite comparable levels of cell-free infection (**Fig. 1b**), reduced ACE2 binding (**Fig. 2c**), and reduced cell-cell fusion (**Fig. 2e**).

## Omicron spike exhibits less S1 shedding

Given the reduced ACE2 binding of Omicron, we speculated that its spike may be less stable. We therefore sought to explore the stability of the Omicron S in parallel with other VOC spikes. First, we examined ACE2-induced S1 subunit shedding by transfecting HEK293T cells with variant S constructs. Cells treated with or without sACE2 and S1 containing-media were collected, and S1 subunit was immune-precipitated using anti-Flag beads. We observed reduced S1 shedding in the Omicron variant for both sACE2-treated and untreated cells (**Fig. 3a**), despite relatively comparable levels of S expression in the cells, with the exception of Delta (**Fig. 3a**). While this may be related to the low ACE2 binding of Omicron variant, distinct mutations unique to Omicron spike, especially those in the furin cleavage site, like contribute to their differential S1 shedding.

**Fig. 3.**
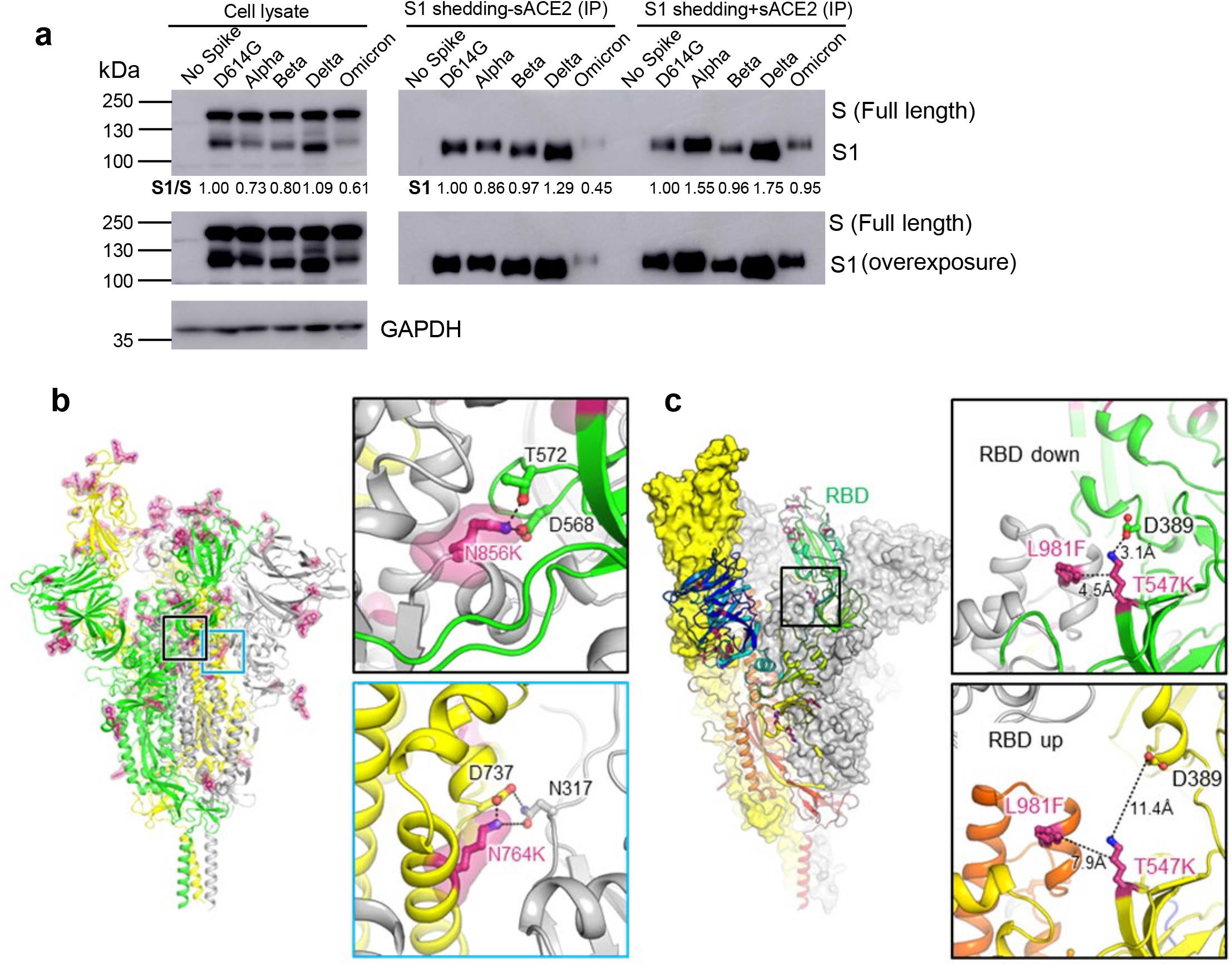
The Omicron S protein exhibits low S1 shedding, consistent with predicted increase in stability. (**a**) HEK293T cells were transfected with S constructs and treated with or without sACE2. Cell lysate and cell culture media were collected after sACE2 treatment, and shed S1 subunit in cell culture media was immune-precipitated with anti-Flag beads. Blots were probed with anti-S1 and anti-GAPDH, and S1 shedding was quantified by NIH ImageJ by setting D614G to 1.0. (**b**) Structure of Omicron spike protein viewed from side. Mutations of Omicron are highlighted by red sticks and semi-transparent red surfaces. RBD of the yellow protomer is in an “up” conformation. Upper inset: the mutation N856K enables formation of salt-bridge and hydrogen bond with the residues D568 and T572, respectively on the neighboring protomer (green). Lower inset: the mutation N764K enables formation of hydrogen bond with residue N317 on the neighboring protomer (grey) and salt-bridge with residue D737 on the same protomer (yellow). (**c**) Omicron mutations T547K and L981F stabilize RBD in “down” conformation. Structure of Omicron spike protein by homology modeling was illustrated as surfaces (yellow and grey protomer) and ribbon (rainbow protomer). Upper inset: When RBD is in “down” conformation, Omicron mutation T547K enables formation of salt-bridge with residue D389 located on the base of RBD. Mutation T547K and L981F together enhance the hydrophobic interaction between the neighboring protomers (green and grey). Lower inset: When RBD is in “up” conformation, the interactions between these residues are disrupted.

To provide additional molecular insights, we created a structural model of the Omicron spike protein using the cryo-EM structure of D614G strain spike (PDB 7KRR) as a homology modeling template (**Fig. 3b**). Structural analysis of the inter-protomer interface indicated an increment in the Omicron spike (6397 Å^2^) compared to the G614 spike (6017.4 Å^2^). The overall solvation free energy gain (ΔiG) shifts from -59.6 kcal/mol to -60.4 kcal/mol, reflecting the enhanced hydrophobic interaction at the Omicron S trimer interface (**Table S1**). Moreover, additional hydrophilic interactions are introduced by Omicron mutations that stabilize the spike trimer formation, including the N856K mutation, which enables formation of a salt-bridge and a hydrogen bond with residues D568 and T572, respectively, on the neighboring protomer. Moreover, the N764K mutation, jointly with the residue D737, forms hydrogen bonds with residue N317 on the neighboring protomer. These inter-protomer interactions enhance association between protomers, as well as between subunits S1 and S2 (**Fig. 3b inset**). The structural model also revealed two critical conformation-stabilizing mutations, T547K and L981F, located close to the base of the spike RBD. When RBD is in the “down” conformation, K547 is in close proximity to both residue 981 in the core of S trimer and residue D389 on the base of RBD (**Fig. 3c**). The hydrophobic interaction formed by K547 and F981, together with the salt-bridge formed by K547 and D389, would lock RBD in the “down” conformation. The “up” conformation of RBD disrupts the interaction among these three residues (**Fig. 3c inset**).

## Discussion

In this work, we examined the efficacy of vaccine-induced immunity against the Omicron variant, along with molecular and virological features of Omicron spike, in parallel with other variants. We found that, while recipients of two-doses of mRNA vaccines showed minimal neutralization of the Omicron variant, recipients of a booster dose, either Moderna or Pfizer, exhibited much stronger neutralizing capacity against Omicron. The fold-difference between nAb titers against D614G and Omicron in the boosted group (3.8-fold) vs. that in the two-dose vaccine group (22.9-fold) revealed that a booster dose not only raises nAb levels, but also increases the breadth of circulating nAbs, a finding that is similar to a recent report by Pfizer^21^. While underlying mechanisms for this remain unclear, enhanced breadth of protection likely is due to additional affinity-maturation following a third antigen exposure. As the Omicron variant may have evolved from a lineage distinct from Delta^22^, it is unsurprising that Omicron exhibited strong resistance to nAb from unvaccinated Delta-infected ICU patients, with Omicron exhibiting ∼8-fold lower nAb sensitivity than Delta for this group.

One surprising finding is that, distinct from Alpha, Beta, and Delta, the Omicron spike exhibited reduced binding to soluble ACE2, which likely accounts for, at least in part, its lower cell-cell fusion efficiency. This may indicate a fitness cost following an accumulation of RBD mutations while under selective pressure for nAb escape. Additionally, reduced cell-cell fusion would reduce cytotoxicity and could contribute to a lower virulence for the Omicron variant, which has been tentatively reported by anecdotal evidence^23^.

Alternatively, reduced ACE2 binding could be reflective of a strategy to reduce premature S inactivation that would enhance virus transmissibility. This possibility is supported by the low S1 shedding observed for Omicron, and confirmed by enhanced stability of the RBD-closed conformation in homology modeling. Notably, infectivity of Omicron in HEK293T-ACE2 cells was not impaired, indicating that it retains functional receptor utilization. However, Omicron exhibited reduced infection in CaLu-3 cells. SARS-CoV-2 is thought to predominantly utilize a TMPRSS2-mediated plasma membrane entry route in CaLu-3 cells, while a cathepsin-B/L-mediated endosomal entry route dominates in HEK293T-ACE2 cells^24,25^. Thus, Omicron S may have evolved to be less fusogenic and utilizes plasma membrane fusion less efficiently in order to minimize potential cytopathogenic effect.

Indeed, the Omicron variant exhibited enhanced cell-to-cell transmission, which would facilitate virus spread. Cell-to-cell transmission is commonly used by many viruses, including SARS-CoV-2, and is a highly efficient mechanism of virus spread within a host^20,26,27^. Enhanced cell-to-cell transmission may help compensate for other observed defects in the Omicron S protein, such as reduced ACE2 binding and fusogenicity. Notably, cell-to-cell transmission of SARS-CoV-2 does not absolutely require ACE2, and extended cell-cell fusion by its spike impairs cell-to-cell transmission^20^. Additionally, cell-to-cell transmission is resistant to neutralizing antibodies, implicating another potential mechanism of Omicron immune evasion^20,26,27^.

Overall, our report highlights the need for booster dose administration to better combat the emerging Omicron variant, and that reformulation of existing mRNA vaccines to target Omicron may not be necessary with a three-dose vaccine regimen. However, the emergence of future divergent variants may further compromise even boosted immunity. Evidence to date suggests no significant enhancement of virulence for Omicron; however, a definitive conclusion will require population level studies as this variant spreads to new locales. Indeed, continued monitoring of emerging SARS-CoV-2 variants is vital to allow for rapid investigation of their transmissibility, neutralization resistance, and pathogenicity.

## Figure Legends

**Figure S1. Neutralization of SARS-CoV-2 Omicron variant by vaccination status and vaccine type**. (**a**) NT_50_ values for HCWs who received two doses of Moderna mRNA-1273 (n = 20) or Pfizer/BioNTech BNT162b2 (n = 28) are plotted by vaccine type. (**b**) NT_50_ values for recipients of Moderna mRNA-1273 (n = 6) or Pfizer/BioNTech BNT162b2 (n = 17) booster doses are plotted by vaccine type. (**c-g**) Post second vaccine dose and post booster dose NT_50_ values are plotted pairwise for HCWs for which both time points were analyzed (n = 18) against the D614G (**c**), Alpha (**d**), Beta (**e**), Delta (**f**), and Omicron (**g**) variants. (**h**) NT_50_ values for unvaccinated (n = 12) and vaccinated (n = 7) Delta-wave COVID-19 ICU-patients are plotted according to vaccination status. Bars in panels a, b and h represent means +/- standard error and statistical significance in all cases was determined by two-tailed t-test with Welch’s correction. Dashed lines indicate the threshold NT_50_, which was set to 80. P-values are represented as **p < 0.01.

**Figure S2. Omicron spike surface expression and sACE2 binding:** (**a-d**) The gating strategy for one representative experiment is shown for determine the single cell population (**a**), the S1 positive population by N-terminal Flag tag (**b**), the S1 positive population by anti-S1 antibody (**c**), and the sACE2 positive population (**d**). (**e**) HEK293T cells were transfected with S expression constructs and stained for FACS with anti-Flag or soluble ACE2-Fc fusion protein (sACE2). (**f**) Mean fluorescence intensity for sACE2 signal was normalized with anti-Flag S1 signal to determine relative ACE2 binding.

## Methods

### Vaccinated and ICU patient cohorts

Vaccinated HCW samples were collected under approved IRB protocols (2020H0228 and 2020H0527). Sera were collected 3-5 weeks post second vaccine dose for 48 HCWs which included 20 Moderna mRNA-1273 and 28 Pfizer/BioNTech BNT162b2 vaccinated HCWs. In the study group, 23 HCWs received homologous vaccine booster doses 34-42 weeks post second dose, these included. Sera was then collected from these 23 HCWs 1-11 weeks post vaccine booster dose. These samples included 6 Moderna mRNA-1273 and 17 Pfizer/BioNTech BNT162b2 boosted HCWs.

D614G-wave ICU and hospitalized non-ICU patient samples were collected under an approved IRB protocol (OSU 2020H0228) as previously described^15^. Delta-wave ICU patient samples were collected under an approved IRB protocol (2020H0175). Plasma samples were collected 3 days after hospitalization. Where detectable, the strain of SARS-CoV-2 infecting the ICU patients was determined by viral RNA extraction on nasal swabs with QIAamp MinElute Virus Spin kit followed by RT-PCR (CDC N1 F: 5′-GACCCCAAAATCAGCGAAAT-3′; CDC N1 R: 5′-TCTGGTTACTGCCAGTTGAATCTG-3′; CDC N2 F: 5′-TTACAAACATTGGCCGCAAA-3′; CDC N2 R: 5′-GCGCGACATTCCGAAGAA-3′) and Sanger sequencing to identify virus strain. Due to risk of infection, Delta-wave ICU patient plasma samples were treated with 1% Triton X100 to inactivate virus. A starting dilution for virus neutralization assays of 1:640 was found to not exhibit Trition X100 mediated cell toxicity.

### Cell lines and maintenance

HEK293T (ATCC CRL-11268, CVCL_1926), HEK293T-ACE2 (BEI NR-52511), and HEK293FT-mCAT-Gluc (a gift from Marc Johnson, University of Missouri) cells were maintained in DMEM (Gibco, 11965-092) supplemented with 10% FBS (Sigma, F1051) and 1% penicillin-streptomycin (HyClone, SV30010). CaLu-3 cells were maintained in EMEM (ATCC, 30-2003) supplemented with 10% FBS and 1% penicillin-streptomycin. All cells were maintained at 37°C and 5% CO_2_.

### Plasmids

We utilized a previously reported pNL4-3-inGluc lentivirus vector which is based on ΔEnv HIV-1 and bears a *Gaussia* luciferase reporter gene that is expressed in virus target cells but not virus producing cells^15,28^. Additionally, SARS-CoV-2 variant spike constructs with N- and C-terminal flag tags were produced and cloned into a pcDNA3.1 vector by GenScript Biotech (Piscataway, NJ) using Kpn I and BamH I restriction enzyme cloning. A pLenti-hACE2-GFP (a gift from Jacob Yount, The Ohio State University) was used transient expression of ACE2. For transient expression of the Tet-Off (tTA) transcription factor, a pQCXIP-Tet-Off expression plasmid was used (a gift from Marc Johnson, University of Missouri).

### Pseudotyped lentivirus production and infectivity

Lentiviral pseudotypes were produced as previously reported^17^. Briefly, HEK293T cells were transfected with pNL4-3-inGluc and spike construct in a 2:1 ratio using polyethylenimine transfection. Virus was harvested 24, 48, and 72 hrs after transfection.

HEK293T-ACE2 or CaLu-3 cells were infected with pseudotyped virus for each strain, produced in parallel. *Gaussia* luciferase activity was assessed 48 hrs after infection by combining cell culture media with *Gaussia* luciferase substrate (0.1M Tris pH 7.4, 0.3M sodium ascorbate, 10 μM coelenterazine). Luminescence was immediately measured by a BioTek Cytation5 plate reader.

### Virus neutralization assay

Pseudotyped lentivirus neutralization assays were performed as previously described^15,17,18,29^. Briefly, HCW serum or ICU patient plasma was 4-fold serially diluted and equal amounts of infectious SARS-CoV-2 variant pseudotyped virus was added to the diluted serum. Final dilutions of 1:80, 1:320, 1:1280, 1:5120, 1:20480, and no serum control for vaccinated HCW samples; while final dilutions of 1:1280, 1:2560, 1:5120, 1:10240, and no serum control were used for ICU patient plasma to avoid Triton X100 toxicity. Virus and serum were incubated for 1 hr at 37°C and then transferred to HEK293T-ACE2 cells for infection. *Gaussia* luciferase activity was determined 48 and 72 hrs after infection by combining 20 μL or cell culture media with 20 μL of *Gaussia* luciferase substrate. Luminescence was immediately measure by a BioTek Cytation5 plate reader. NT_50_ values were determined by least-squares-fit, non-linear regression in GraphPad Prism 5 (San Diego, CA). Heat maps with NT_50_ generated by GraphPad Prism 9.

### Spike binding sACE2 and anti-Flag and detection by flow cytometry

Virus producing HEK293T cells were harvested 72 hrs after transfection. Cells were dissociated by incubation at 37°C in PBS + 5mM EDTA for 30 min. Then, cells were fixed in 4% formaldehyde in PBS, and stained with sACE2-humanFc fusion protein (construct is a gift of Jason McLellan, University of Texas at Austin) or mouse anti-Flag antibody (Sigma, F3165). Cells were then stained with anti-human-IgG-FITC (Sigma, F9512) or anti-mouse-IgG-FITC (Sigma, F0257) and processed by a Life Technologies Attune NxT flow cytometer. Results were processed using FlowJo v7.6.5 (Ashland, OR). Relative ACE2 binding was then determined by dividing sACE2 signal by anti-Flag signal measured by mean fluorescence intensity.

### Syncytia formation assay

HEK293T cells were co-transfected with a plasmid encoding GFP along with that a variant spike of interest. The transfected HEK293T cells were co-cultured with HEK293T-ACE2 cells 24 hrs after transfection. Cells were imaged 24 hrs after co-culture at 4x magnification on a Leica DMi8 confocal microscope. Representative images were selected.

### Tet-off-based cell-cell fusion assay

Donor HEK293T cells were transfected with an S construct of interest and pQCXIP-Tet-Off in a 1:1 ratio. Target HEK293FT-mCAT-Gluc cells stably expressing tetracycline-responsive element (TRE)-driven *Gaussia* luciferase were transfected with pLenti-hACE2-GFP. 24 hrs after transfection, HEK293FT-mCAT-Gluc and HEK293T cells were dissociated with PBS-EDTA, and cocultured at a 1:1 ratio. Fusion between S and Tet-Off expressing HEK293T cells and ACE2 expressing HEK293FT-mCAT-Gluc cells would drive expression of *Gaussia* luciferase. Thus, at 24 hrs and 48 hrs after co-culture, cell culture media were sampled and assayed for *Gaussia* luciferase activity.

### Spike incorporation into pseudotyped virus

Pseudotyped virus particles were purified by ultracentrifugation through a 20% sucrose cushion. Virus was resuspended in SDS-PAGE loading buffer. Cell lysate from virus producer cells was collected by 30 min incubation of cells on ice in RIPA lysis buffer (50 mM Tris pH 7.5, 150 mM NaCl, 1 mM EDTA, Nonidet P-40, 0.1% SDS) supplemented with protease inhibitor (Sigma, P8340). Samples were run on a 10% acrylamide SDS-PAGE gel and transferred to a PVDF membrane. Membranes were probed with anti-Flag (Sigma, F3165), anti-p24 (Abcam, ab63917; NIH ARP-1513), anti-S1 (Sino Biological, 40150-T62), and anit-β-actin (Sigma, A1978). Anti-mouse-IgG-Peroxidase (Sigma, A5278) and anti-rabbit-IgG-HRP (Sigma, A9169) were used as secondary antibodies where appropriate. Blots were imaged with Immobilon Crescendo Western HRP substrate (Millipore, WBLUR0500) on a GE Amersham Imager 600.

### S1 shedding

HEK293T cells were transfected with S expression constructs. Then, 24 hrs after transfection, cells were treated with or without sACE2 (25 μg/mL) for 4 hrs at 37°C. Cell lysate and cell culture media was harvested. S1-containig cell culture media was incubated with 10 μL of anti-Flag beads (Sigma, F2426) to precipitate S1 subunit. Following immune-precipitation, cell lysate and shed S1 were run on 10% SDS-PAGE gel, transferred, and probed with anti-S1 (Sino Biological, 40150-T62) and anit-GAPDH (Santa Cruz, sc-47724). Anti-mouse-IgG-Peroxidase (Sigma, A5278) and anti-rabbit-IgG-HRP (Sigma, A9169) were used as secondary antibodies.

### Structural modeling

Homology modeling of Omicron spike protein was conducted on SWISS-MODEL server with cryo-EM structure of SARS-CoV2 G614 strain spike (PDB 7KRR) as template. The resulting homo-trimer spike structure has one RBD in up conformation and the other two RBD in down conformation. Residue examination and renumbering were carried out manually with program Coot.

### Molecular contact analysis

Inter-protomer interaction analysis was performed with PDBePISA server. The Omicron spike inter-protomer interface was compared to that of G614 strain in both overall assembly and individual residue levels. Intra-protomer contacts of Omicron mutants were examined with the programs PyMOL and chimeraX. All structural figures were generated with PyMOL. The detail analysis results were listed in Table S1 and Supplementary Table 1. Homology modeling was performed in PyMOL using the cryo-EM structure of D614G strain spike (PDB 7KRR) as homology modeling template to predict Omicron spike protein structure.

### Statistics

Comparisons between multiple groups were made using a one-way ANOVA with Bonferroni post-test. Comparisons between two-groups were made using a two-tailed student’s t-test with Welch’s correction.

## Acknowledgements

We thank Jason Mclellan, David Derse, Marc Johnson, and Ali Ellebedy for providing plasmids, cells, and antibodies. We also thank the NIH AIDS Reagent Program and BEI Resources for supplying important reagents that made this work possible. We thank the Clinical Research Center/Center for Clinical Research Management of The Ohio State University Wexner Medical Center and The Ohio State University College of Medicine in Columbus, Ohio, specifically Francesca Madiai, Dina McGowan, Breona Edwards, Evan Long, and Trina Wemlinger, for logistics, collection and processing of samples. In addition, we thank Sarah Karow, Madison So, Preston So, Daniela Farkas, and Finny Johns in the clinical trials team of The Ohio State University for sample collection and other supports.

## Author Contributions

S.-L.L conceived and directed the project. C.Z., J.P.E., and P.Q. contributed the majority of the experimental work, data processing, and drafting of the manuscript. J.F. performed infectivity assays and provided valuable discussion. K.X. and T.Z. performed homology modeling. C.C., J.S.B., G.L., R.M., R.J.G. provided clinical samples. J.P.E, C. Z., and S.-L. L wrote the paper. P.M. facilitated shipping of the Omicron construct. Y.-M.Z, L.J.S., E.M.O., P.M., and R.J.G. provided insightful discussion and revision of the manuscript.

