## Supplemental Figure 1 for "Neutralization and Stability of SARS-CoV-2 Omicron Variant"

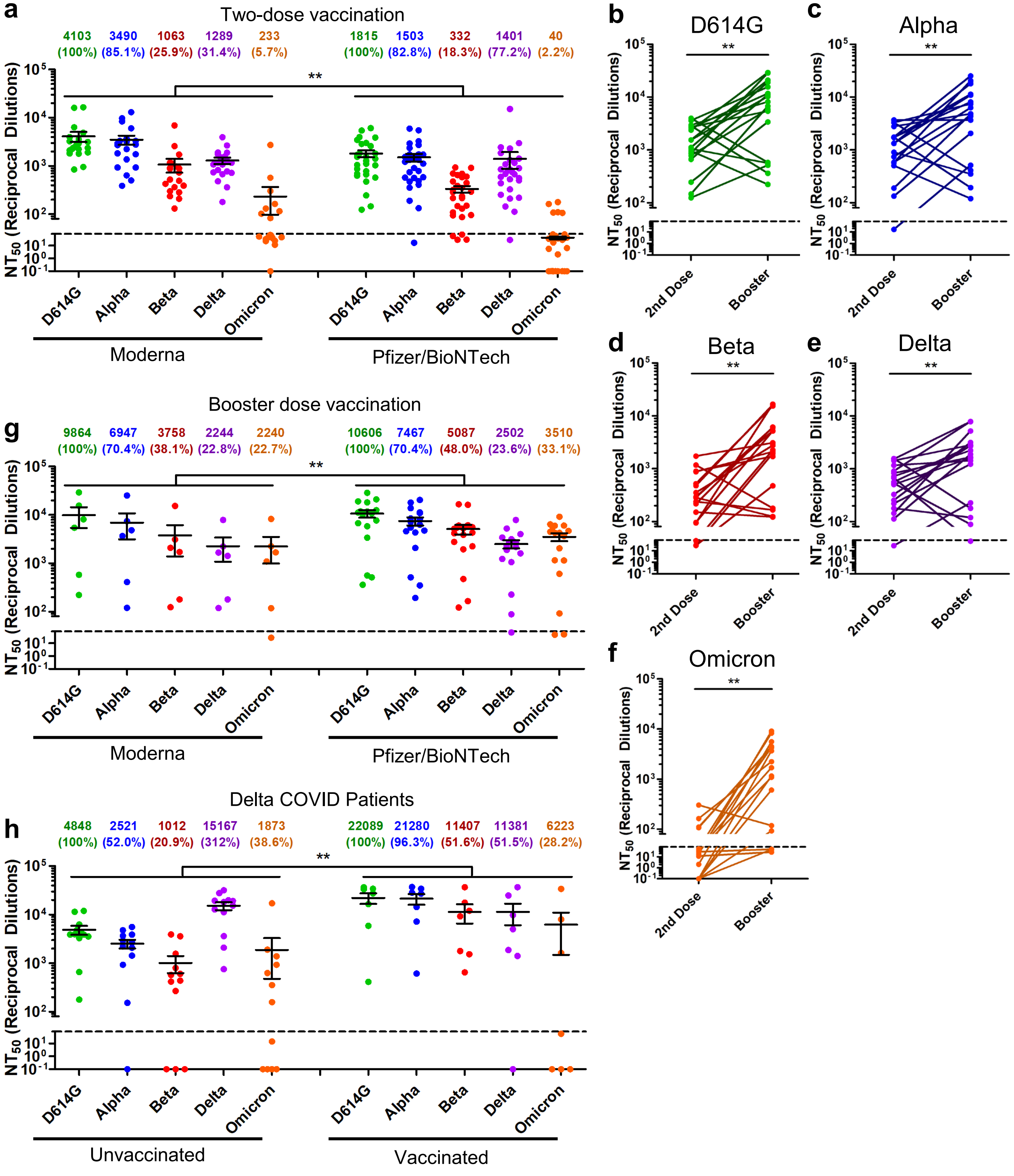


**Figure S1. Neutralization of SARS-CoV-2 Omicron variant by vaccination status and vaccine type.** (**a**) $NT_{50}$ values for HCWs who received two doses of Moderna mRNA-1273 (n = 20) or Pfizer/BioNTech BNT162b2 (n = 28) are plotted by vaccine type. (**b**) $NT_{50}$ values for recipients of Moderna mRNA-1273 (n = 6) or Pfizer/BioNTech BNT162b2 (n = 17) booster doses are plotted by vaccine type. (**c-g**) Post second vaccine dose and post booster dose $NT_{50}$ values are plotted pairwise for HCWs for which both time points were analyzed (n = 18) against the D614G (**c**), Alpha (**d**), Beta (**e**), Delta (**f**), and Omicron (**g**) variants. (**h**) $NT_{50}$ values for unvaccinated (n = 12) and vaccinated (n = 7) Delta-wave COVID-19 ICU-patients are plotted according to vaccination status. Bars in panels a, b and h represent means +/- standard error and statistical significance in all cases was determined by two-tailed t-test with Welch's correction. Dashed lines indicate the threshold $NT_{50}$, which was set to 80. P-values are represented as **p < 0.01.

Supplementary Figure 1
