## Supplemental Figure 2 for "Neutralization and Stability of SARS-CoV-2 Omicron Variant"

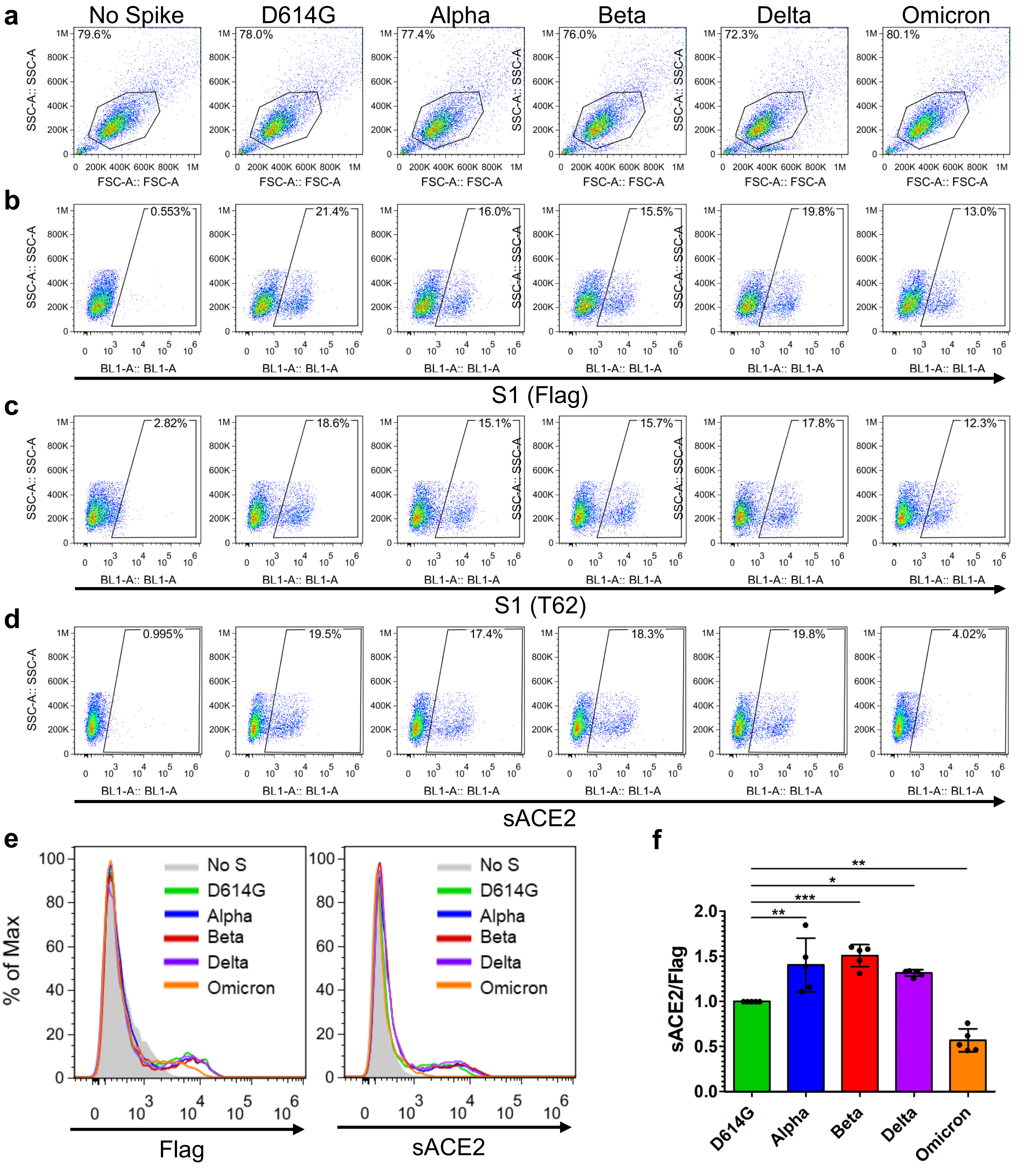


**Figure S2. Omicron spike surface expression and sACE2 binding:** (**a-d**) The gating strategy for one representative experiment is shown for determine the single cell population (**a**), the S1 positive population by N-terminal Flag tag (**b**), the S1 positive population by anti-S1 antibody (**c**), and the sACE2 positive population (**d**). (**e**) HEK293T cells were transfected with S expression constructs and stained for FACS with anti-Flag or soluble ACE2-Fc fusion protein (sACE2). (**f**) Mean fluorescence intensity for sACE2 signal was normalized with anti-Flag S1 signal to determine relative ACE2 binding.

Supplementary Figure 2
