## Supplemental Table 1 for "Neutralization and Stability of SARS-CoV-2 Omicron Variant"

### Table S1. Omicron spike trimer interface analysis by PDBePISA

#### a Overall interface comparison between Omicron and G614 S trimer

| | Protomer1 | | | Protomer2 | | | interface | ΔiG | ΔiG | #HB | #SB |
|---|---|---|---|---|---|---|---|---|---|---|---|
| | iNat | iNres | Surface Å² | iNat | iNres | Surface Å² | area, Å² | kcal/mol | P-value | | |
| Omicron | 660 | 188 | 59124 | 714 | 191 | 58896 | 6397.6 | -60.4 | 0.092 | 56 | 14 |
| G614 | 623 | 176 | 56231 | 670 | 181 | 56824 | 6017.4 | -59.6 | 0.056 | 46 | 10 |

Note:
iNat: the number of interfacing atoms.
iNres: the number of interfacing residues.
ΔiG: Solvation energy effect, kcal/mol
HB: Hydrogen bond
SB: Salt bridge

#### b Unique hydrophilic interaction contributed by Omicron mutations

| Hydrogen bonds | | | Salt bridges | | |
|---|---|---|---|---|---|
| Protomer 1 | Dist. [Å] | Protomer 2 | Protomer 1 | Dist. [Å] | Protomer 2 |
| A:ASN 317[ OD1] | 3.79 | B:LYS 764[ NZ ] | A:ASP 568[ OD2 | 2.80 | B:LYS 856[ NZ ] |
| A:ASP 568[ OD2] | 2.80 | B:LYS 856[ NZ ] | | | |
| A:THR 572[ OG1] | 2.79 | B:LYS 856[ NZ ] | | | |

#### c Individual interface contact contributed by Omicron mutations

| | HSDC | ASA | BSA | ΔiG |
|---|---|---|---|---|
| A:ASN 417 | | 55.99 | 8.14 \|\| | -0.12 |
| A:LYS 547 | | 117.56 | 65.99 \|\|\|\|\|\| | 0.31 |
| A:GLY 614 | H | 75.10 | 57.25 \|\|\|\|\|\|\|\| | 0.04 |
| A:LYS 969 | H | 95.26 | 26.12 \|\|\| | 0.29 |
| B:LYS 764 | H | 77.78 | 11.22 \|\| | -0.41 |
| B:TYR 796 | H | 192.06 | 50.52 \|\|\| | 0.24 |
| B:PHE 981 | | 124.99 | 44.63 \|\|\|\| | 0.41 |

ASA: Accessible Surface Area, Å²
BSA: Buried Surface Area, Å²
ΔiG: Solvation energy effect, kcal/mol
||||: Buried area percentage, one bar per 10%
